# The developmental stage of the medulloblastoma cell-of-origin restricts Hedgehog pathway usage and drug sensitivity

**DOI:** 10.1101/2021.01.22.427790

**Authors:** Marlinde J. Smit, Inna Armandari, Irena Bockaj, Tosca E.I. Martini, Walderik W. Zomerman, Zillah Siragna, Tiny G.J. Meeuwsen, Frank J.G. Scherpen, Mirthe H. Schoots, Martha Ritsema, Wilfred F.A. den Dunnen, Eelco W. Hoving, Judith T.M.L. Paridaen, Gerald de Haan, Victor Guryev, Sophia W.M. Bruggeman

**Author notes:** Corresponding author. Correspondence (S.B.). These authors contributed equally (Inna Armandari, Irena Bockaj, Tosca E.I. Martini).

## Abstract

Sonic Hedgehog (SHH) medulloblastoma originates from the cerebellar granule neuron progenitor (CGNP) lineage that depends on Hedgehog signaling for its perinatal expansion. While SHH tumors exhibit overall deregulation of this pathway, they also show patient age-specific aberrations.

To investigate if the developmental stage of the CGNP can account for these age-specific lesions, we analyzed developing murine CGNP transcriptomes and observed highly dynamic gene expression as function of age. Cross-species comparison with human SHH medulloblastoma showed partial maintenance of these expression patterns, and highlighted low primary cilium expression as hallmark of infant medulloblastoma and early embryonic CGNPs. This coincided with reduced responsiveness to upstream Shh pathway component Smoothened, while sensitivity to downstream components Sufu and Gli was retained.

Together, these findings can explain the preference for *SUFU* mutations in infant medulloblastoma and suggest that drugs targeting the downstream SHH pathway will be most appropriate for infant patients.

## Introduction

Medulloblastoma is a malignant tumor of the cerebellum that frequently affects children. It consists of four main transcriptional subgroups that can be further subdivided upon additional molecular profiling [1–5]. The Sonic Hedgehog (SHH) subgroup of medulloblastoma, which accounts for thirty percent of all medulloblastoma cases, is believed to originate from the cerebellar granule neuron progenitor (CGNP) population that critically depends on Sonic Hedgehog pathway signaling for its perinatal expansion [4–11]. Sonic Hedgehog is a secreted morphogen that controls development and patterning in many organs including the central nervous system [12,13]. The pathway becomes active when SHH binds to its receptor PATCHED (PTCH1), which relieves inhibition of SMOOTHENED (SMO) and induces MYCN [12–15]. Activated SMO inhibits tumor suppressor protein SUFU and stimulates processing of the GLI factors into transcriptional activators. In the absence of the SHH morphogen, transcription of SHH target genes is actively repressed by SUFU mediated formation of GLI repressors or direct transcriptional repression [16–18].

In line, SHH medulloblastoma is characterized by an overall deregulation of SHH signaling that is often accompanied by mutually exclusive mutations in SHH pathway components, underlining the importance of this pathway in driving tumorigenesis [1,19]. Interestingly, there is also patient age-related heterogeneity within this subgroup, suggesting that developmental factors affect tumor biology [2,3,19–23]. For instance, infant and adult medulloblastoma display distinct gene expression patterns, as well as differences in copy number alterations and tumor localization. Strikingly, mutations in SHH pathway genes are also correlated with patient age [19]. Whereas *PTCH1* mutations occur across all age groups, *SUFU* mutations are almost exclusively found in infant patients; GLI2, NMYC and TP53 mutations in older children; and *SMO* mutations in adults.

These latter findings raise the question why the pathway is differentially perturbed depending on patient age, as these mutations would presumably have identical outcomes, *i*.*e*., enhanced activation of SHH target genes. One explanation is that the tumor cells-of-origin undergo changes in sensitivity to, or usage of, the Sonic Hedgehog signaling pathway during development, which would then provoke age-specific oncogenic lesions in the pathway [24]. The presumed cell-of-origin for SHH medulloblastoma is part of the CGNP lineage, a highly dynamic cell population that spans over three weeks of mouse and two years of human development [25], with the precise moment of transformation remaining under debate [6,11,34,26–33]. In mice, future CGNPs become specified in the upper rhombic lip (uRL) of the hindbrain around embryonic day E13.5, from where they migrate across the surface of the cerebellar primordium [26–29]. Here, they form a secondary germinal zone termed external granular layer (EGL), with a peak in proliferation occurring around birth that is driven by Purkinje neuron secreted Shh. Once they reach maturity, terminally differentiating granule neurons cease proliferation and migrate inwards until they reach their final destination in the internal granular layer (IGL) of the cerebellum [9–11,28,30]. Besides the longer gestational period, the development of human CGNPs generally resembles that of the mouse [31].

To study the potential impact of the CGNP developmental age on medulloblastoma outcome, we have taken advantage of a transgenic mouse model that allows the prospective isolation of the entire CGNP cell lineage during embryonic and postnatal development [35,36]. As reported before, we confirmed that CGNPs exhibit dynamic changes in gene expression in time, highlighting the identity changes of the CGNP population as neural development progresses [4,5,35]. We also confirmed that CGNPs resemble human SHH medulloblastoma and importantly, that early embryonic CGNPs co-segregate with the youngest patients, corroborating a linear relationship between cell-of-origin and patient age. In particular, we found that primary cilium expression was low across young CGNPs and patients, which was somewhat unexpected given the importance of primary cilia for Shh pathway activity [37–46]. In line, early embryonic CGNPs displayed a partial unresponsiveness to Smo-mediated (*i*.*e*., upstream) pathway stimulation, preventing increased proliferation. They were however sensitive to deletion of *Sufu* or inhibition of Gli, both downstream Shh components, which in contrast to Smo are known to have cilium-independent functions [47,48]. These observations may have clinical implications, as patients with early developmental medulloblastoma are likely to benefit from drugs targeting the downstream, primary cilium-independent part of the SHH pathway [49].

## Results

### Developing cerebellar granule neuron progenitors undergo dynamic changes in the expression of genes implicated in medulloblastoma

To address if intrinsic changes in the developing medulloblastoma cell-of-origin could contribute to the age-dependent mutations in SHH pathway components found in SHH medulloblastoma, a transgenic mouse model was employed that allows lineage tracing and prospective isolation of the developing murine CGNP cell lineage from their specification in the cerebellar primordium onwards (Figure 1A) [29,35,50]. Accordingly, we observed that treatment of pregnant transgenic mice with a single dose of tamoxifen at E13.5 induced acute and stable labeling of CGNPs with tdTomato in the offspring (Figure 1B). We dissected fluorescent cerebella at E15.5, E17.5, P0 (day of birth), P7 (postnatal day 7), P14 and P30, purified tdTomato expressing CGNPs and generated transcriptomes (Figure 1A,C-E and Table S1). Subsequent principal component analysis revealed that more than 70% of all variation in gene expression between samples could be explained by developmental age, demonstrating that CGNPs indeed undergo significant changes in gene expression as a function of time (Figure 1C).

**Figure 1.**
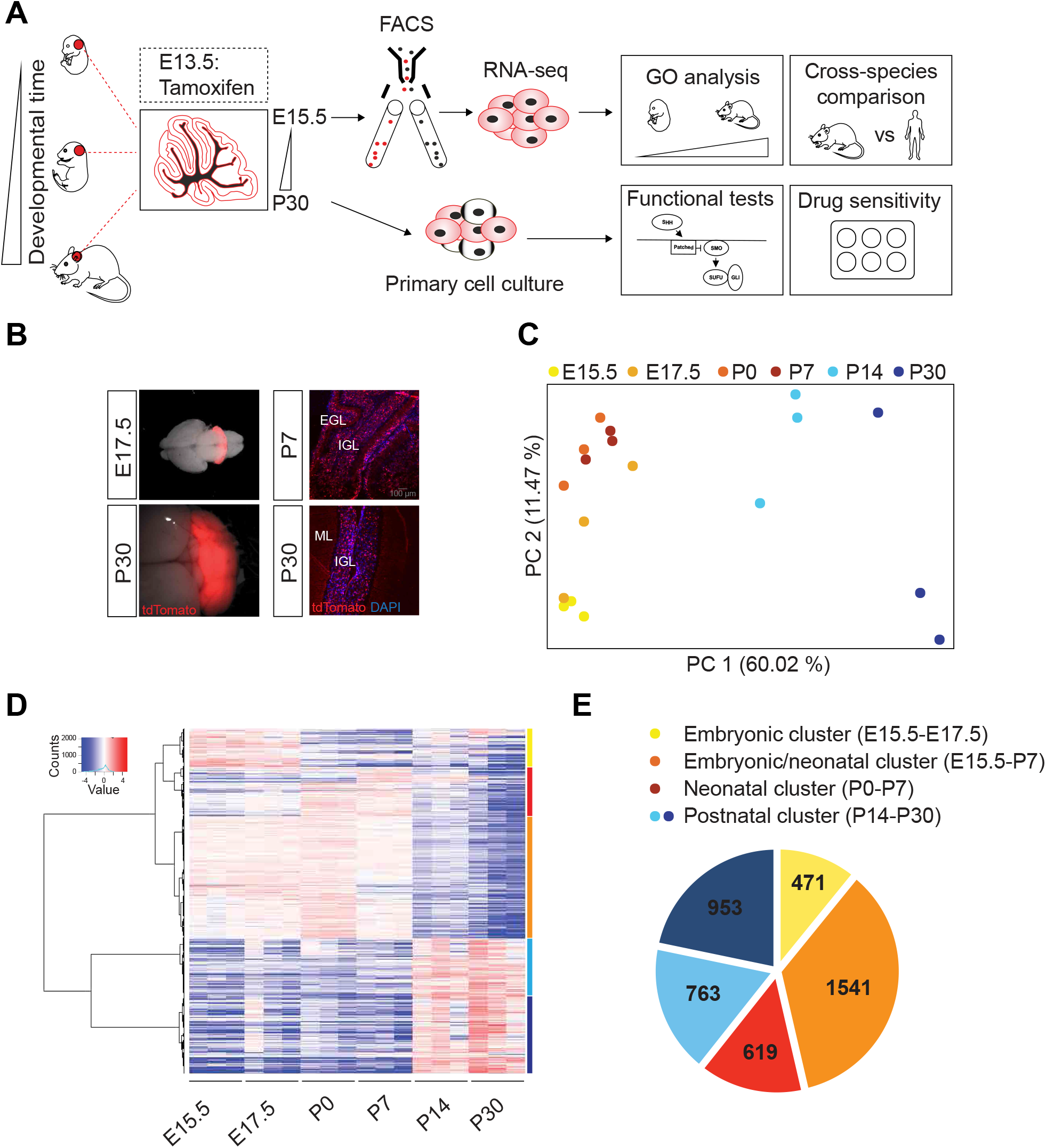
Developing cerebellar granule neuron progenitors (CGNPs) exhibit age-specific gene expression. (A) Schematic overview of the experimental workflow. Embryonic CGNPs are labeled with tdTomato following a single Tamoxifen pulse administered to pregnant females. Fluorescent cerebella are dissected at different time points between E15.5 and P30. tdTomato^+^ cells are sorted by FACS for further experiments. Note: for cell culture experiments, CGNPs are treated with 4-hydroxytamoxifen in vitro (E=embryonic day; P=postnatal day, FACS=fluorescence-activated cell sorting). (B) Stereoscopic (left panels) or microscopic images (right panels) showing direct tdTomato fluorescence in whole mount cerebella and sagittal sections, respectively (EGL=external granular layer; IGL=internal granular layer; ML=molecular layer). (C) Principal component analysis of E15.5, E17.5, P0, P7, P14 and P30 CGNP transcriptomes (three biological replicates per developmental time point. For embryonic time points, CGNPs from n=4 embryos were pooled per sample; for postnatal time points, individual mice were analyzed). (D) Heatmap showing differentially expressed CGNP genes as function of developmental time point (unsupervised hierarchical clustering analysis). Genes are clustered according to the branching of the clustering tree into five major clusters: yellow cluster (E15.5-E17.5); orange cluster (E15.5-P7); red cluster (P0-P7); light blue and dark blue clusters (P14-P30). (E) Pie chart summarizing five major clusters of differentially expressed genes. Yellow cluster (E15.5 - E17.5, n=471 genes); orange cluster (E15.5 - P7, n=1541 genes); red cluster (P0-P7, n=619 genes); light blue cluster (P14 - P30, n=763); and dark blue cluster (P14 - P30, n=943).

To further explore these dynamic changes, we performed unsupervised hierarchical clustering analysis to identify genes that were differentially expressed (DE) between at least two time points (Figure 1D and Table S2). We next divided the DE genes into five major gene clusters according to the clustering tree. The yellow cluster contained genes that were highest expressed at embryonic time points (E15.5 and E17.5, n=471 genes), the orange cluster genes that were high during embryonic and early postnatal time points (E15.5-P7, n=1541 genes), genes in the red cluster were highest expressed at early postnatal time points (P0 and P7, n=619 genes), and light and dark blue clusters contained the genes that were induced upon granule neuron maturation (P14 and P30, n=763 and 943 genes, respectively) (Figure 1D,E). To identify enriched biological processes (Gene Ontology) within these gene clusters, we used the Database for Annotation, Visualization and Integrated Discovery (DAVID) and visualized the data using Cytoscape (Figures 2 and S1, and Table S3) [51]. At the early stages of CGNP development (yellow cluster), there is a strong enrichment of processes associated with early neural development such as axon and dendrite formation and developmental transcription factor-driven gene expression, but also glycolysis (Figures 2 and S1). In comparison, processes enriched throughout embryonic and neonatal stages (orange cluster) are biased towards cell proliferation and mitosis. This surge in proliferation may be accompanied by increased DNA damage, as also processes related to the DNA damage response (DDR) are frequent in this cluster. The end of cerebellar proliferation is heralded by the appearance of cell cycle arrest processes, which are specific to early neonatal CGNPs (red cluster). Of note, the vast majority of orange cluster genes show the highest relative gene expression levels at P0, suggesting that proliferation of CGNPs born around E13.5 peaks around birth and declines at P7 (Figure 1D). Two orange cluster processes (GO terms: covalent chromatin modification and ATP-dependent chromatin remodeling) stand out, as genes involved in chromatin regulation like *Arid2*, are known to be mutated in medulloblastoma [52,53]. During the final stages of CGNP differentiation (light and dark blue clusters), processes related to neuronal connectivity, differentiation and regeneration are enriched, as well as major signal transduction routes like Bmp, TGF-beta and PI3K signaling that are also known to play a role in medulloblastoma [22,54,55].

**Figure 2.**
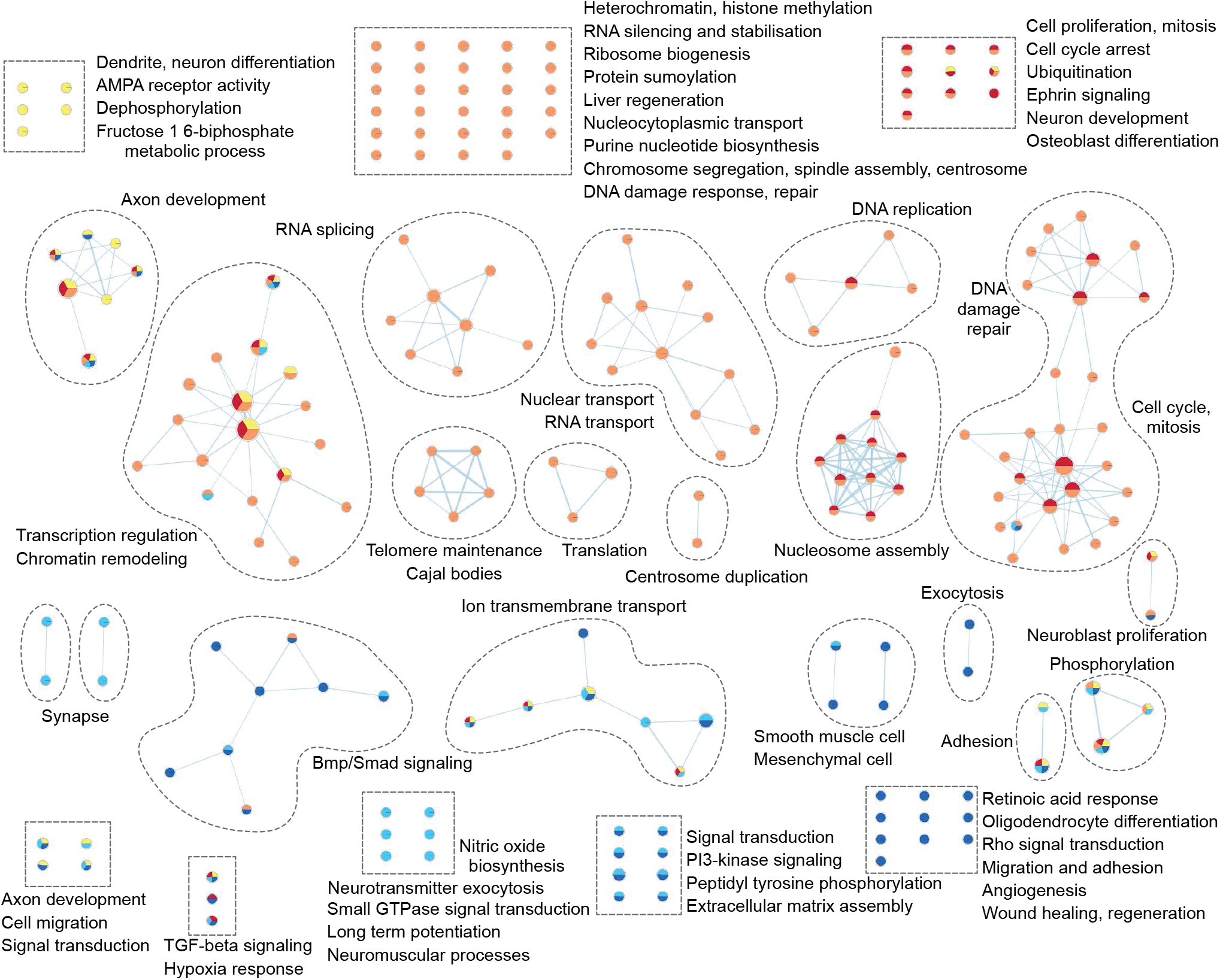
Specific biological processes are enriched during CGNP development. Gene ontological analysis shows enriched biological processes in the different CGNP age-specific gene clusters. Each node represents a biological process. Related biological processes are grouped and labeled by biological theme (curved dashed lines). Individual biological processes are assembled in rectangular boxes (dashed lines). Biological processes connected by edges have genes in common. Enriched biological processes were determined with the Database of Annotation, Visualization and Integrated Discovery (DAVID), v.6.8 (Benjamini-corrected q=0.1, p=0.01) and visualized with the Enrichment Map app in Cytoscape. Yellow nodes: E15.5 - E17.5 cluster; orange nodes: E15.5 - P7 cluster; red nodes: P0 - P7 cluster; light blue nodes: P14 - P30 cluster; dark blue nodes: P14 - P30 cluster (See also Figure S1).

To further explore a potential link between temporal CGNP gene expression and medulloblastoma, we extracted the individual gene expression profiles of n=40 genes commonly mutated in medulloblastoma and grouped them as suggested by Northcott et al (Figures 3A and S2) [3]. We found striking peaks in the expression of the genes involved in cell cycle and genome maintenance at P0, and genes involved in chromatin and transcription regulation at P7, indicating that the age of the cell-of-origin could be related to mutations found in medulloblastoma. Interestingly, we observed the highest induction of *TP53* expression at birth, which is most frequently mutated in children over the age of three [19].

**Figure 3.**
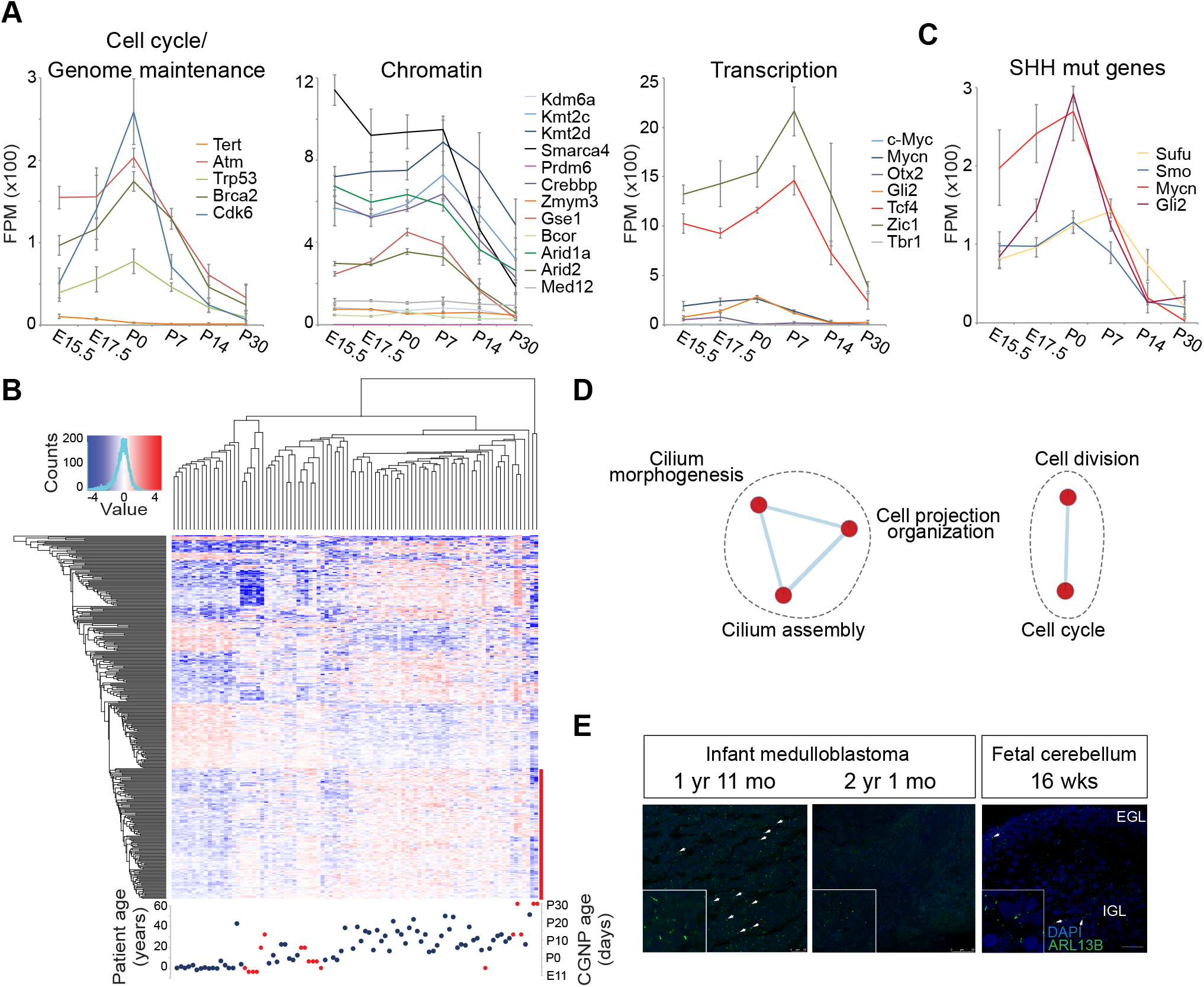
Cell cycle regulation and primary cilia biogenesis are age-dependent processes in CGNPs and medulloblastoma. (A) Gene expression profiles of CGNP genes commonly mutated in medulloblastoma, extracted from the RNA-seq data set. Curves represent the average expression level from three biological replicates, error bars indicate standard deviation (FPM=fragments per million, E=embryonic day, P=postnatal day) (See also figure S2). (B) Cross-species comparison. Heatmap showing unsupervised hierarchical clustering of differentially expressed human medulloblastoma genes (patient-age groups: 0-3 years, 4-11 years, 12 years and older) and CGNP orthologous genes. Blue dots represent patient samples, red dots represent CGNP samples. Lowest gene cluster indicated by red bar. (C) Gene expression profiles of CGNP genes associated with the Sonic Hedgehog signaling pathway, extracted from the RNA-seq data set. (D) Enrichment map representing biological processes enriched in the lower (red) gene cluster of the cross-species comparison heatmap (b). No enriched processes were found in the upper gene clusters. Each node represents a biological process. Related biological processes are grouped and labeled by biological theme (curved dashed lines). Biological processes connected by edges have genes in common. Enriched biological processes were determined with the Database of Annotation, Visualization and Integrated Discovery (DAVID), v.6.8 (Benjamini-corrected q=0.1, p=0.01) and visualized with the Enrichment Map app in Cytoscape. (E) Confocal images showing ARL13B protein expression (primary cilium marker) in infant medulloblastoma (left and middle panels) and human fetal cerebellum (right panel). White arrows indicate primary cilia (EGL=external granular layer; IGL=internal granular layer). Scale bars, 25 µm.

### Age-specific CGNP gene expression is preserved in human medulloblastoma

To investigate if overall CGNP gene expression is reflected in human SHH medulloblastoma, we performed an independent cross-species comparison between an existing cohort of human SHH medulloblastoma and our murine CGNP samples (Figure 3B) [19]. Hereto, we selected the human genes that were differentially expressed between SHH medulloblastoma patients of three age groups, *i*.*e*., 0-3 years of age (infants/toddlers), 4-11 years of age (children), and 12 years and older (older children/adults). In agreement with earlier publications, we confirmed that infant and adult medulloblastoma have distinct gene expression patterns, with childhood medulloblastoma forming an intermediate group [19,20]. Interestingly, unsupervised hierarchical clustering analysis of these human genes together with their murine counterparts showed that embryonic mouse CGNPs co-cluster with the infant medulloblastomas, and older CGNPs with tumors from older patients (Figure 3B). This demonstrates that CGNPs resemble human medulloblastoma, and that the age of the patient is partially reflected by the age of the medulloblastoma cell-of-origin.

### Cell cycle regulation and primary cilium biogenesis are age-related processes in CGNPs and medulloblastoma

We next re-examined the CGNP transcriptome data with the purpose of finding an explanation for the existence of age-specific Hedgehog pathway mutations. We hereby hypothesized that these genes are differentially susceptible to oncogenic mutation as a function of time, because they are not equally important at all stages of development. If true, there could be differences in peak gene expression levels for the mutated Hedgehog pathway genes as we had observed for *Tp53*. To test this, we extracted the individual gene expression profiles from genes associated with the Hedgehog signaling pathway from our CGNP transcriptome data. However, while we observed that Shh target genes like *Gli1/2, Ccnd1/2* and *Ptch1/2*, and to a lesser extent *Mycn*, had clear expression peaks at P0, this was not evident for either the *Sufu* or *Smo* genes that exhibit striking age-specific mutation patterns (Figures 3C and S2). Thus, only *Gli2 and Mycn*, which are predominantly mutated in children over the age of three in conjunction with *TP53*, show age-specific gene expression peaks [19].

We subsequently searched for alternative age-related processes that could impose differential Hedgehog pathway usage on the CGNPs. Hereto, we subjected the gene clusters from the cross-species comparison to gene ontological analysis (Figure 3B,D and Table S4). We identified only five enriched biological processes, which were all enriched in the older CGNP/patient group (lower gene cluster, indicated by red bar). These processes were either involved in cell proliferation or in primary cilium formation (Figure 3D and Table S5). Since it is known that primary cilia are required for SHH signaling, it seemed paradoxical that younger patients have relatively low primary cilia gene expression. We stained a small panel of infant medulloblastoma (n=5) for ARL13B, a marker for primary cilia [56], and confirmed that there is large variation in the number of ciliated cells between different patients, with some tumors hardly expressing any primary cilia (Figure 3E, left and middle panels) [57,58]. Thus, low primary cilium expression can occur in infant medulloblastoma.

We next investigated primary cilium expression in normal developing CGNPs. In human second trimester fetal cerebellum, we found ciliated cells in both EGL and IGL, the latter being most prominent (Figure 3E, right panel). As we had no access to first trimester embryonic human cerebellum that harbors the early specified uRL and presumptive EGL cells, we subsequently turned to the developing murine cerebellum to establish the dynamics of primary cilium expression from the uRL stage onwards. In contrast to the general view that most cells have a primary cilium, we found that in the E12.5-E13.5 mouse cerebellum, cells in the uRL and future EGL are largely devoid of primary cilia whereas surrounding brain structures are ciliated (Figure 4A) [43,59]. As development progresses, the number of ciliated cells in the EGL and later also IGL, increases (Figure 4B-D). We noticed that at neonatal stages, ciliated CGNPs were more abundant towards the outer EGL in agreement with earlier studies [45,46]. However, ciliated CGNPs were most frequent in the IGL, and these cilia were also significantly longer (Figure 4B-D). Altogether, this demonstrates that both primary cilium expression and length are dynamic during CGNP development, and that at early phases of medulloblastoma cell-of-origin development, primary cilium expression is rare.

**Figure 4.**
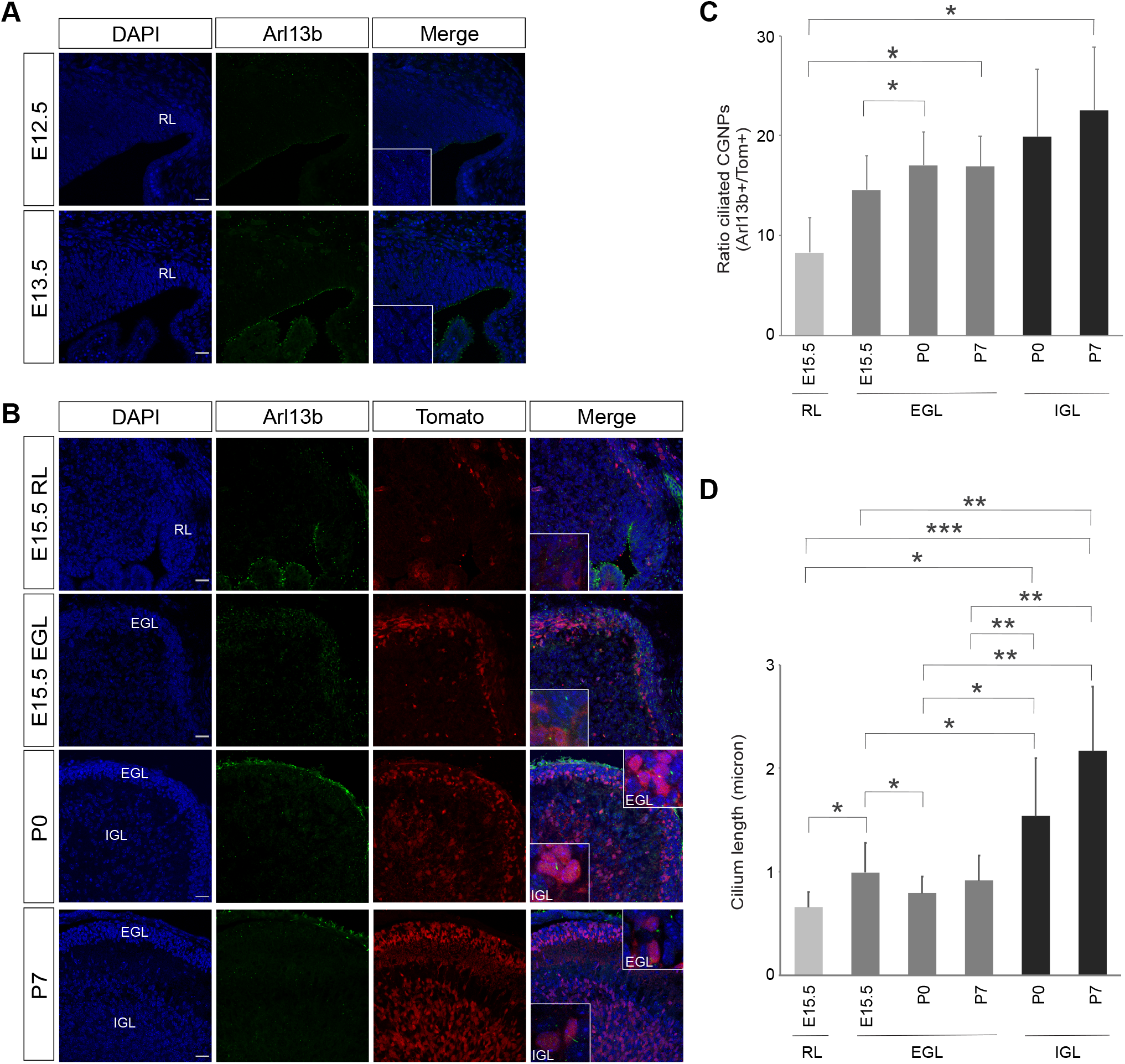
Primary cilium expression and length in the developing murine cerebellum. (A) Confocal images showing Arl13b expression in developing mouse cerebellum (E12.5 and E13.5) (RL=rhombic lip; E=embryonic day). Scale bars, 25 µm. (B) Confocal images showing Arl13b and tdTomato expression in the developing mouse cerebellum (E15.5 EGL, E15.5 IGL, P0, P7) (P=postnatal day, EGL, external granular layer; IGL, internal granular layer). Insets showing higher magnification. (C) Chart showing the average ratio of ciliated CGNPs (*i*.*e*., Arl13b^+^/tdTomato^+^ cells) per location and developmental timepoint. Three independent biological replicates were measured. Error bars indicate standard deviation. p values were determined using a one-sided paired t-test. * p < 0.05. (D) Chart showing the average primary cilium length per location and developmental timepoint. Three independent biological replicates were measured. Error bars indicate standard deviation. p values were determined using a one-sided paired t-test. * p < 0.05; ** p < 0.01; *** p < 0.001.

### Sufu and Smoothened expression is dynamic and partially non-overlapping during cerebellar development

Both *Smo* and *Sufu* function is essential for Hedgehog signaling in the cerebellum, with Smo acting upstream in relaying the signals from Shh morphogen-bound Ptch, and Sufu acting downstream in receiving signals from activated Smo to cease the inhibition of Gli [60,61]. A major difference though is their dependence on the primary cilium for pathway activation. Smo requires the primary cilium for its function, whereas Sufu is also known to have cilium-independent activity [37,38,40,42,47,48,62]. Hence, if a subset of infant medulloblastoma is derived from the earliest specified CGNPs, this could explain the increased frequency of *SUFU* mutations, as these cells are mostly non-ciliated. We therefore checked if during cerebellar development, Sufu and Smo proteins were expressed in patterns consistent with this hypothesis (Figure 5). In general, Sufu exhibited the broadest expression (Figure 5A). At embryonic day E15.5, Sufu was most prominently expressed in the presumptive EGL, and was also found in the uRL. At postnatal stages, Sufu expression became more restricted towards the outer EGL and upper layers of the IGL. In striking contrast, Smo was absent from the uRL or (presumptive) EGL, and appeared restricted to the area where the IGL forms (Figure 5B).

**Figure 5.**
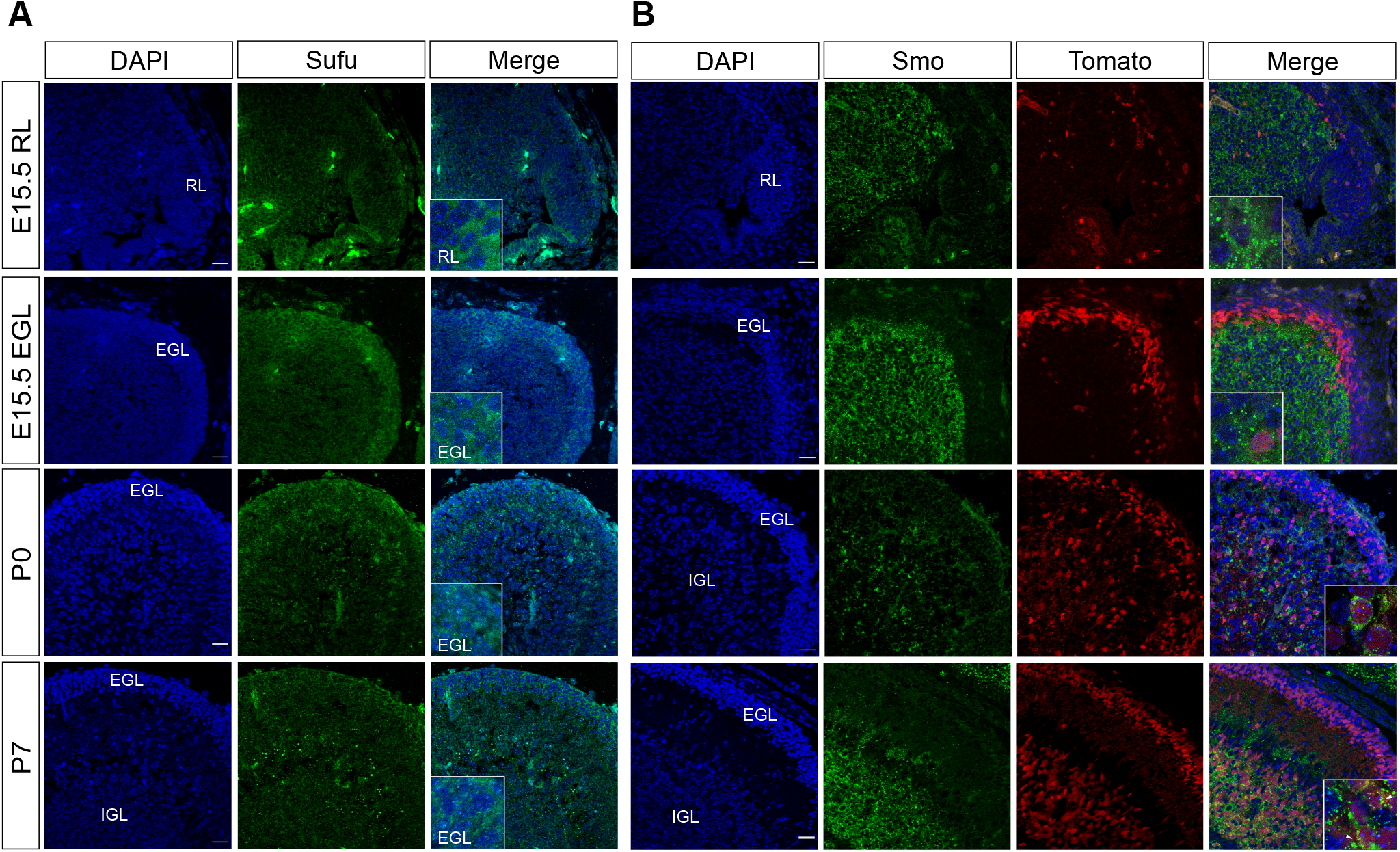
Differential Sufu and Smoothened expression patterns in the developing cerebellum. Confocal images showing (A) Sufu and (B) Smoothened (Smo) protein expression in the E15.5 RL, E15.5 EGL, P0 and P7 cerebellum. CGNPs were identified by tdTomato staining. Counterstaining by DAPI (RL=rhombic lip; EGL=external granular layer; IGL=internal granular layer; E=embryonic day; P=postnatal day). Insets showing higher magnification. White arrow heads indicate primary cilia. Scale bars, 25 µm.

### Embryonic CGNPs have reduced sensitivity to Smoothened manipulation but can be activated by downstream Shh signaling

We then set out to functionally test if embryonic CGNPs are differentially sensitive to Shh pathway component manipulation compared to early postnatal CGNPs, as suggested by the differential expression of Sufu, Smo, and primary cilia. Hereto, we isolated *Math1-CreER*^*T2*^; *tdTomato* CGNPs from either E15.5 or P7 cerebellum and cultured them for brief periods of time to preserve their primary status (48-72 hrs) [64–67]. We observed ciliated CGNPs in both cultures with P7 CGNPs having longer cilia similar to the *in situ* stainings, suggesting that cilia are not completely absent from E15.5 CGNPs (Figure 6A). In agreement with earlier publications, we found that upon overexpression of oncogenic *Smo* (SmoM2) or knockdown of *Sufu*, P7 CGNPs increase proliferation, indicating engagement of upstream and downstream Shh pathway segments at this time (Figures 6B,C and S3A,C,D) [9,11,60]. However, in E15.5 CGNPs, only *Sufu* knockdown cells showed increased proliferation, whereas SmoM2 overexpression had no effect (Figure 6B,C).

**Figure 6.**
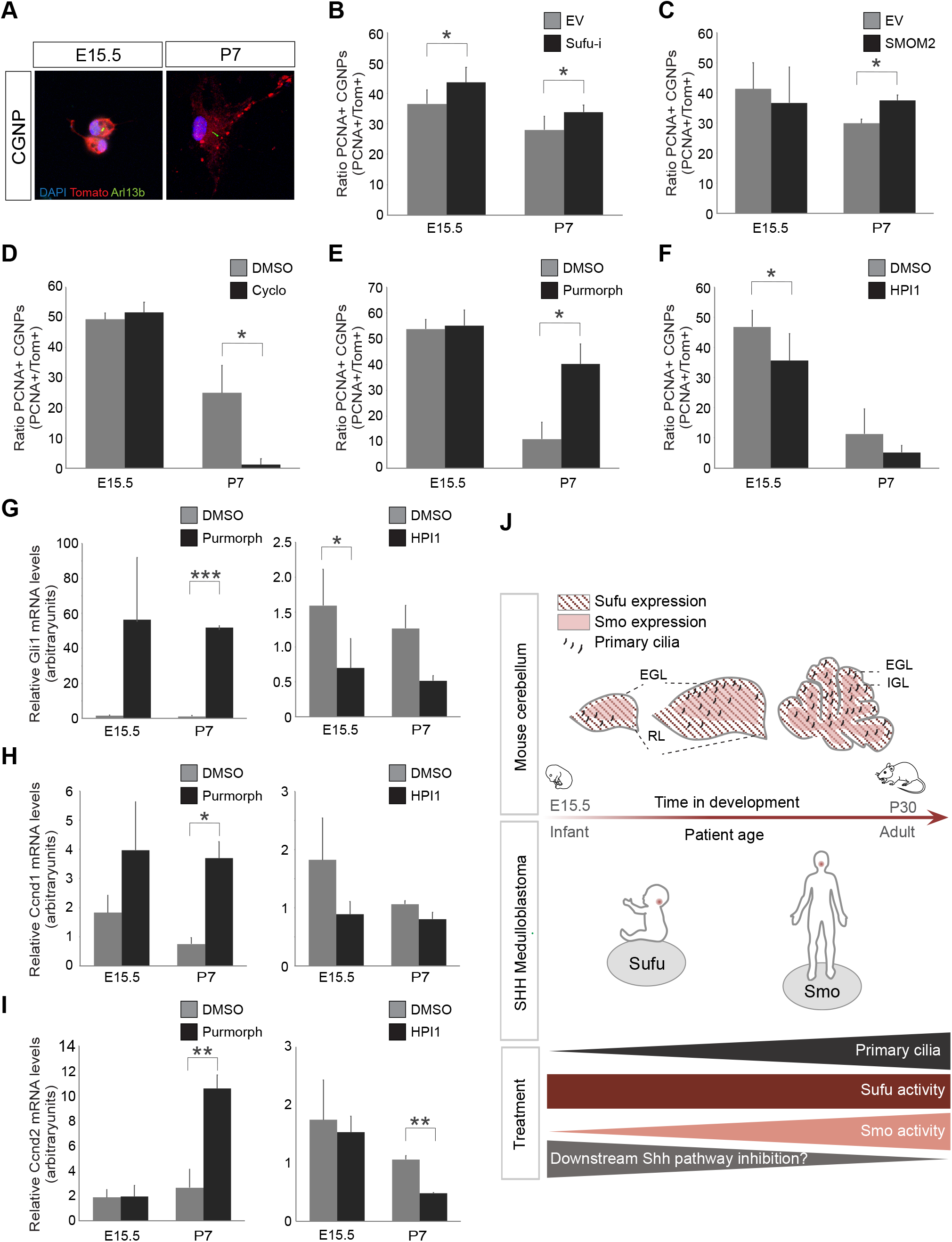
Embryonic CGNPs are insensitive to Smoothened manipulation. (A) Confocal images showing Arl13b and tdTomato expression in E15.5 and P7 primary CGNP cultures. Counterstaining by DAPI. (B) Chart showing the average ratio of PCNA^+^/tdTomato^+^ cells after Sufu-i shRNA or empty vector (EV) control retroviral transduction, in E15.5 (left bars) or P7 CGNP primary cultures (right bars), from three biological replicates. Error bars indicate standard deviation. p values were determined using a one-sided two sample t-test. * p < 0.05. See also Figure S3. (C) Chart showing the average ratio of PCNA^+^/tdTomato^+^ cells after SMOM2 or empty vector (EV) control retroviral transduction, in E15.5 (left bars) or P7 CGNP primary cultures (right bars), from two or three biological replicates. Error bars indicate standard deviation. p values were determined using a one-sided two sample t-test. * p < 0.05. See also Figure S3. (D-F) Charts showing the average ratio of PCNA^+^/tdTomato^+^ cells after treatment with DMSO (control) or 5 µM Cyclopamine (d), 250 nM Purmorphamine (e), or 2.5 µM HPI1, in E15.5 (left bars) or P7 CGNP primary cultures (right bars), from at least three biological replicates. Error bars indicate standard deviation. p values were determined using a one-sided paired t-test. * p < 0.05. See also Figure S3. (G-I) Charts showing relative Gli1 (g), Cyclin D1 (h), or Cyclin D2 (i) mRNA expression levels compared to Gapdh in Purmorphamine (left panels) or HPI1 (right panels) treated E15.5 and P7 CGNPs, as determined by qRT-PCR from at least three biological replicates. Error bars indicate standard deviation. p values were determined using a two-sided paired t-test. * p < 0.05, ** p < 0.01, *** p < 0.001. See also Figure S3. (J) Proposed model. Sonic Hedgehog (SHH) medulloblastoma derives from the cerebellar granule neuron progenitor (CGNP) population. CGNPs are specified in the rhombic lip (RL) of the early embryonic cerebellum, from where they migrate across the cerebellar surface to form the external granular layer (EGL). Upon terminal differentiation, CGNPs migrate inwards to form the definitive internal granular layer (IGL). Expression and length of primary cilia, which are important components of upstream SHH signaling, increases as CGNP development progresses. Thus, if oncogenic transformation takes place at late stages of CGNP development, *SMO* is preferentially mutated as the primary cilia enhance oncogenic SMO activity. However, if CGNP oncogenic transformation occurs at an early embryonic stage, *SUFU* is preferentially mutated, which controls downstream Sonic Hedgehog signaling independently from the primary cilium. This implies that targeted therapy for infant SHH medulloblastoma should be directed towards downstream tumor-driving mechanisms (E=embryonic day, P=postnatal day).

Prompted by these results, we wanted to address if E15.5 CGNPs are sensitive to treatment with Smo inhibitor Cyclopamine, derivatives of which are being used in the treatment of relapsed SHH medulloblastoma [68]. Whereas proliferation of P7 CGNPs was strongly diminished by Cyclopamine treatment as was shown before, E15.5 CGNPs were almost completely insensitive (Figure 6D and S3B). Likewise, treatment with Purmorphamine, a Smo agonist, had no effect on E15.5 CGNP proliferation whereas P7 CGNPs were highly sensitive (Figure 6E). In contrast, when we treated E15.5 CGNPs with HPI1, a compound that inhibits the downstream Shh target Gli1, we did see a significant reduction in proliferation, implying that the downstream Shh pathway is functional in early embryonic CGNPs (Figure 6F). To further address this at the molecular level, we analyzed the expression of key Shh target genes (e.g., *Gli1, Patched, Nmyc, Cyclin D1* and *Cyclin D2*) in Purmorphamine or HPI1 treated E15.5 and P7 CGNPs (Figures 6G-I and S3E,F). Surprisingly, whereas E15.5 CGNPs did not increase proliferation upon Purmorphamine treatment, they showed significant induction of *Gli1* gene expression, suggesting that these early cells are not completely unresponsive to upstream Shh signaling. However, in line with the absence of a proliferative response, they did not increase expression of *Cyclin D1* and Cyclin *D2* like P7 CGNPs (Figure 6H,I). Together, these data show that the age of the medulloblastoma cell-of-origin determines the extent of the response to Shh pathway activation, and as a consequence has an impact on SHH drug sensitivity (Figure 6J).

## Discussion

Whereas Sonic Hedgehog medulloblastoma is characterized by an overall deregulation of the Hedgehog signaling pathway, there are also abnormalities specific to patient age [3,19,20,32,69]. In this study, we set out to address if intrinsic changes in developmental processes between young and older cerebellar granule neuron progenitors, a cell lineage known to be highly susceptible to SHH medulloblastoma formation, can account for the age-specific characteristics of this tumor type [6,36].

### The developmental age of the medulloblastoma cell-of-origin is partially reflected in tumors

The idea that naturally occurring changes in identity of the tumor cell lineage-of-origin impact on tumor outcome, is supported by studies showing differences in tumor onset and phenotype in relation to timing of tumor induction, although the type of genetic lesion and intra-tumoral heterogeneity are also expected play a role [4,5,32,70–73]. Indeed, when we compared gene expression patterns between developing CGNPs, we found unique enriched biological processes at each developmental stage, and several of those have been linked to SHH medulloblastoma. A linear chronological relationship between CGNP and patient age was evident in some but not all cases [2,3,19,20,69]. For instance, neurotransmission activity and neural development are enriched in both early embryonic CGNPs and infant medulloblastoma [2]. PI3K signaling on the other hand, also associated with infant medulloblastoma, is mostly enriched in the late postnatal CGNP samples representing maturing granule neurons [19].

Intriguingly, there was a strong enrichment for processes related to cell cycle control and the DNA damage response throughout early CGNP development with a noticeable peak around birth. As these processes are overrepresented within the older SHH medulloblastoma subtypes, this suggests that the SHH-induced perinatal surge in CGNP proliferation, possibly coinciding with increased DNA damage due to replication stress, is a critical event during cerebellar development that brings along the risk of developing child- and adulthood medulloblastoma [2–5]. The strong increase in *TP53* expression at birth is particularly interesting, as it might be related to the high incidence of *TP53* mutations and occurrence of chromothripsis in children [21].

### Differential Hedgehog pathway regulation during cerebellar development might impact on infant SHH medulloblastoma outcome

An important remaining question is the precise nature of the cell-of-origin for infant SHH medulloblastoma. In our cross-species comparison, we found that the infant SHH medulloblastomas clustered towards the earliest embryonic CGNPs, suggesting an early embryonic granule neuron progenitor (up to E15.5) as cell-of-origin [32]. This is in line with elegant single cell sequencing studies, which showed that a subset of SHH MB clustered with early embryonic cerebellar cells that were identified by the authors as CGNP and unipolar brush cell precursors of comparable age [4,5]. However, initially we were puzzled by the fact that both infant SHH tumors and early embryonic CGNPs were characterized by low primary cilium expression, as the latter is believed to be indispensable for SHH signaling [37,44,74]. Because how can we reconcile early embryonic CGNPs being cell-of-origin for infant medulloblastoma exhibiting deregulated Shh signaling, if these cells are not capable of Shh signaling? And moreover, how can we then explain that there is a higher incidence of *SUFU* mutations, and complete lack of *SMO* mutations, in these tumors [19]?

We now think that Sufu’s unique role in the SHH pathway might be the answer to these questions. It had already been shown that in contrast to *Smo, Sufu* does not absolutely depend on the primary cilium to exert its control over Hedgehog target gene expression [41,42,47,48]. In addition, another vital difference between *Smo* and *Sufu* is their effect on Hedgehog signaling, as Smo activates, and Sufu represses target gene expression [12– 15,18]. We therefore favor a model in which transformation of early stage CGNPs into infant medulloblastoma requires *SUFU* deletion, as this would induce precocious pathway activation independent of a Hedgehog signal; but not SMO activation that in the absence of primary cilia would not have a proliferative effect [58]. This idea is supported by studies from others showing a significantly earlier role in cerebellar development for *Sufu* compared to *Smo* and *Shh* [30,60,61,75–78]. Furthermore, mutant cerebella lacking primary cilia also show relatively late developmental phenotypes, in line with primary cilia being most important during perinatal Shh-induced CGNP proliferation [62,79]. Whether the presence or absence of the primary cilium itself plays a functional role in this, awaits further investigation.

### Implications for future medulloblastoma research

Only recently, it was demonstrated that the four consensus subgroups of medulloblastoma can be further sub-classified using a combination of molecular profiling techniques [2,3,19,20,80]. These studies have provided a wealth of information on the molecular genetics, as well as putative regions of origin for these tumors, which are both essential pieces of information for developing accurate preclinical models and subsequential drug testing. For instance, while several mouse models for medulloblastoma have been generated, none of them has faithfully recapitulated infant SHH medulloblastoma, suggesting that the correct cell-of-origin has not been properly targeted [81]. The importance of *bona fide* preclinical modeling is underscored by our finding that early embryonic CGNPs, the putative cells-of-origin for infant SHH medulloblastoma, are inherently insensitive to Smo inhibition.

Thus, in addition to taking into account the level at which the Hedgehog pathway is compromised when designing targeted therapy, intrinsic characteristics of the cell-of-origin preserved in the tumor should also be taken into consideration [82,83]. Especially in infants, who typically do not receive radiotherapy, it is crucial to use drugs precisely tailored to their specific medulloblastoma subtype [84].

## Materials and methods

### Experimental animals

The Math1-CreER^T2^; tdTomato compound transgenic mouse strain was derived from the Math1CreER^T2^ and Ai14 mouse strains (The Jackson Laboratory), and in a C57BL6/mixed background. Mice were conventionally housed, fed *ad libitum*, and routinely genotyped by PCR. Timed matings were performed overnight, with the following morning considered E0.5. Pregnancies were detected by measuring female weight gain at E13.5. A subset of pregnant females received a single dose of Tamoxifen (2 mg/100 μl peanut oil, Sigma) by oral gavaging at E13.5. Pregnant females were killed by asphyxiation (CO_2_), neonatal mice until the age of P7 were killed by decapitation. Offspring from different gender were randomly assigned. All animal experiments were approved by the Institutional Animal Care and Use Committee of the University Medical Center Groningen, the Netherlands.

### CGNP single cell suspension preparation

For transcriptional analyses, CGNPs were harvested from E15.5, E17.5, P0, P7, P14, and P30 tdTomato^+^ dissected cerebella from offspring from Tamoxifen-treated females. For primary cell cultures, CGNPs were harvested from E15.5 and P7 days old cerebella from offspring of non-treated females. Cerebellar dissection was performed using a stereomicroscope. E15-P30 Cerebella were dissociated with a Papain dissociation kit following the manufacturer’s instructions (Worthington). Following papain treatment, ovomucoid was added to stop the reaction. For P14 and P30 cerebella, an additional Percoll gradient step was performed to remove myelin and debris. Cell suspensions were filtered through a 40 μm cell strainer prior to further processing.

### Primary CGNP cell cultures

For embryonic CGNP cultures (E15.5), n=10-20 dissected cerebella were pooled. For P7 CGNP cultures, n=2-4 dissected cerebella were pooled. Single cell suspensions were prepared as described above. Cells were pelleted and resuspended in the appropriate culture media. For E15.5 CGNPs: BME media supplemented with 1% N2, 2% B27 (Invitrogen), and 1 μM 4-Hydroxytamoxifen (Sigma). For P7 CGNPs: DMEM-F12 media supplemented with 1% N2, 1.5% sucrose, 5 μm HEPES (Invitrogen), 0,25 μg/ml Shh (R&D systems), and 1 μM 4-Hydroxytamoxifen. For proliferation assays and RNA isolation, 250.000 E15.5 CGNPs or 500.000 P7 CGNPs cells were seeded into poly-D-Lysine (100 μg/ml, Sigma) coated 24 wells plates (E15.5) or 12 wells plates (P7). For confocal imaging, 100.000 E15.5 CGNPs or 60.000 P7 CGNPs were seeded onto Ibidi 8-chamber glass slides coated with poly-D-Lysine.

For drug treatments, E15.5 or P7 CGNPs were plated as described above in the presence 5 μM Cyclopamine (Merck), 250 nM Purmorphamine (Stem Cell Technologies), 2.5 μM HPI1 (Tocris), or DMSO, and incubated for 48 hrs. Cells were seeded as one or more (if sufficient cells were available) technical replicates per experiment, and at least three biological replicates (*e*.*g*., samples derived from different individual mice) were tested.

### Lentiviral transductions

The Sufu shRNA construct was generated by cloning a Sufu-targeting 22-mer oligonucleotide into a modified pRRL-SFFV-IRES-GFP plasmid (restriction sites XhoI and EcoRI), with tNGFR replaced by GFP (kind gift from Dr. Hein schepers) (See also Table S6) [85].

293T producer cells were transfected with prrl-SFFV-IRES-GFP, prrl-SFFV-IRES-GFP-Sufu or SmoM2 (W535L)-pcw107-V5 (Addgene) using Fugene HD transfection reagent (Promega). Appropriate CGNP culture media was added to the producer cells 16 hours prior to virus harvest. CGNPs were subsequently incubated with lentiviruses for 2 hours, after which virus was gradually replaced with normal culture media to prevent cell death. Cells were fixed at 72 hours following transduction.

### Quantification of proliferating or ciliated cells

For quantification of PCNA positive CGNPs in primary cultures, random microscopic images (n=10 per sample) were taken and processed using CellProfiler 3.1.9 software (Broad Institute, cellprofiler.org) using an adapted Counting and Scoring pipeline. Average ratios of double positive cells were calculated and tested for significance (p<0.05) using paired t-tests. For primary cilium quantification, Arl13b and tdTomato (double) positive cells were counted per developmental time point in the RL, EGL and IGL in fluorescence microscopic images (n=3 per age group), with the use of Fiji software. Average ratios of double positive cells were determined and tested for significance using paired t-tests.

Cilium length was measured manually using Fiji. We took the average (from n=12 cilia per sample, except for the E15.5 samples where less cilia were present) of the cilium length in RL, EGL and IGL per developmental time point (n=3 per age groups) and tested for significance using paired t-tests.

### Immunofluorescence

CGNPs were fixed with 100% MeOH (for PCNA staining) or 4% formaldehyde (for Arl13b), and blocked with 0.1% triton, 1% BSA and 0.05% Tween in PBS. Primary antibodies were: Arl13b (Proteintech, 17711-1-AP, 1:400), PCNA (Abcam, ab29, 1:1000) and RFP (Rockland, 600-401-379, 1:500). Secondary antibodies were Alexa Fluor 488 (1:500) and Alexa Fluor 568 (1:500) (Invitrogen). Cells were counterstained with DAPI (Sigma).

Brains from Tamoxifen-treated Math1-CreER^T2^; tdTomato mice were dissected from the skull, fixed with 4% formaldehyde, and cryoprotected with a sucrose gradient (10%, 20 % and 30% sucrose in PBS). They were embedded into Tissue-Tek O.C.T. compound (Sakura-FineTek) and snap frozen in liquid nitrogen. Human fetal brain and medulloblastoma tissue was obtained from the University Medical Center Groningen. Local ethics committee approval was granted for use of the patient material. Cryosections (10 μm) were generated on a Leica cryostat. Antigen retrieval was performed by boiling tissue sections in Citrate buffer (100 mM, pH 6.0), except for Arl13b and mCherry. Sections were blocked with 5% normal goat serum and 0.1% Triton (Cell Signaling) in PBS. Primary antibodies were: Arl13b (Proteintech, 17711-1-AP, 1:100), mCherry (SICGEN, AB0040-200, 1:200), PCNA (Abcam, ab29, 1:1000), RFP (Rockland, 600-401-379, 1:500), Sufu (Abcam, ab28083, 1:50), Smo (Abcam, ab38686, 1:500), Histone H3S10ph (Active Motif, 39636, 1:100). Secondary antibodies were Alexa Fluor 488 (1:500) and Alexa Fluor 568 (1:500) (Invitrogen). Slides were counterstained with DAPI (Sigma), and mounted with Vectashield (Vector Laboratories).

### Microscopy

Whole mount images of fluorescent embryonic and postnatal mouse brain were made on an Olympus stereozoom SZX-16 microscope. Primary cells were imaged on an EVOS FL inverted fluorescence microscope (Life technologies). Fluorescent tissue sections were imaged on a Leica TCS SP8 confocal microscope, and analyzed using Fiji [86].

### Quantitative RT-PCR

RNA from E15.5 and P7 CGNPs was isolated using the RNeasy Mini Kit (Qiagen). cDNA was synthesized using random hexamer primers, dNTPs and Ribolock (Thermo Scientific).

Reverse transcription was performed using Reverse Transcriptase (Thermo Scientific). Quantitative RT-PCR was performed on a BioRad CFX connect Real-time system using universal SYBR® Green supermix (BioRad). Relative gene expression was calculated using the 2^-ΔΔ^CT method. Expression levels were normalized to housekeeping gene Gapdh. For primers, see Table S6.

### CGNP FACS and RNA isolation for RNA-Seq

Cerebellar single cell suspensions (E15.5-P30) were subjected to fluorescence-activated cell sorting (FACS) for tdTomato^+^ cells, using a MoFlo Astrios cell sorter (Beckman Coulter). tdTomato^+^ cells (CGNPs) were collected in 1.5 ml tubes, centrifuged, and snap frozen. For the E15.5 and E17.5 time points, tdTomato positive cerebella from n=4-8 embryos from the same litter were pooled prior to FACS to constitute biological replicate; for P0-P30 time points, individual mice were sorted to constitute a biological replicate.

RNA was extracted using the NucleoSpin RNA XS kit (Macherey-Nagel), following the manufacturer’s instructions. RNA concentration was measured with a QuBit RNA HS assay kit (Thermo Fisher) and analyzed on a bioanalyzer using the Agilent RNA 6000 Pico Kit (Agilent). 2.5-10 ng of RNA per sample was used as input for library preparation.

### Library preparation and RNA-Seq

For library prepping, the Clontech SMARTer stranded total RNA-seq kit-pico was used according to the manufacturer’s instructions for mammalian sample input. Three biological replicates were sequenced for each time point (Illumina HiSeq 2500). On average, 16.7 million reads (63 bp single-end) were generated for each replicate. Reads were aligned to the mouse reference genome (GRCm38 assembly, gene annotation from Ensembl release 84, http://www.ensembl.org) and quantified using STAR 2.5.3a [87].

### Hierarchical clustering analysis

For hierarchical clustering analysis, differentially expressed genes were called for all possible pairwise comparisons of developmental stages (3 replicates each) using the edgeR package. Genes were selected that significantly changed their expression (FDR<1e-5) in one or more comparisons. Log2 ratios were calculated relative to the average FPM expression of a gene. Unsupervised hierarchical clustering and heatmap plotting was performed using gplots library. The Canberra distance and Ward clustering method was used.

### Cross species comparison

For comparison of gene expression profiles between mouse CGNP and human medulloblastoma patients, a published human medulloblastoma data set was downloaded from GEO (https://www.ncbi.nlm.nih.gov/geo; accession number GSE49243). Expression values were transformed into log2 fold change (compared to average expression of a gene across all patients). Unambiguous orthologs (one to one orthology) were determined using the Ensembl Biomart tool (http://www.ensembl.org/biomart). Unsupervised hierarchical clustering and heatmap plotting was done using gplots library. The Euclidean distance and Average clustering method was used.

### Gene Ontology

Pathway enrichment analysis was performed by uploading lists of differentially expressed genes (gene clusters), to the Database for Annotation, Visualization and Integrated Discovery v6.8 (DAVID), and subsequent analysis for gene ontology of biological processes was performed [88]. Lists of biological processes were imported into the Enrichmentmap app in Cytoscape v3.2.1 [89,90]. Biological processes were Benjamini-corrected with a moderately permissive q value of <0.1 and a p-value of <0.01. Enrichment maps represent biological processes enriched in the gene clusters. Each node represents a biological process grouped and labeled by biological theme. Biological processes connected by edges have genes in common using a Jaccard and Overlap coefficient combined with a similarity cutoff value of 0.375.

## Data and software availability

The accession number for the CGNP RNA-Seq data set is E-MTAB-7399 (https://www.ebi.ac.uk/arrayexpress/).

## Acknowledgements

We are grateful to Dr. Ilia Vainchtein for assistance on cerebellar granule neuron cell sorting, and Dr. Seka Lazare for assistance with RNA-seq. This work was funded by a De Cock Stichting grant to M.S.; an Indonesian Endowment Fund for Education (LPDP) grant (20151022024475) to I.A.; a Rosalind Franklin fellowship from the University of Groningen and European Union to J.P.; a Mouse Clinic for Cancer and Ageing/Large Infrastructure grant from the Netherlands Organisation for Scientific Research (NWO) to G.H., a Stichting Vrienden Beatrix Kinderziekenhuis grant, a Rosalind Franklin fellowship from the University of Groningen, and a KWF Cancer Research career award (RUG 2014-6903) to S.B.

## Author contributions

Conceptualization, M.S. and S.B.; methodology, M.S., J.P., V.G., and S.B.; software and formal analysis, W.Z., M.S., S.B., and V.G.; investigation, M.S., I.A., I.B., T.M., W.Z., Z.S., T.M., F.S., M.R., and S.B.; resources, M.Sc, W.D., J.P., E.H., and G.H.; data curation, V.G.; writing, M.S. and S.B.; writing-review and editing, M.S., J.P., I.B., W.Z., and S.B.; visualization, M.S., W.Z., V.G. and S.B.; supervision and project administration, S.B.; funding acquisition, M.S. and S.B.

## Supplemental information

**Figure S1 related to Figure 2.**
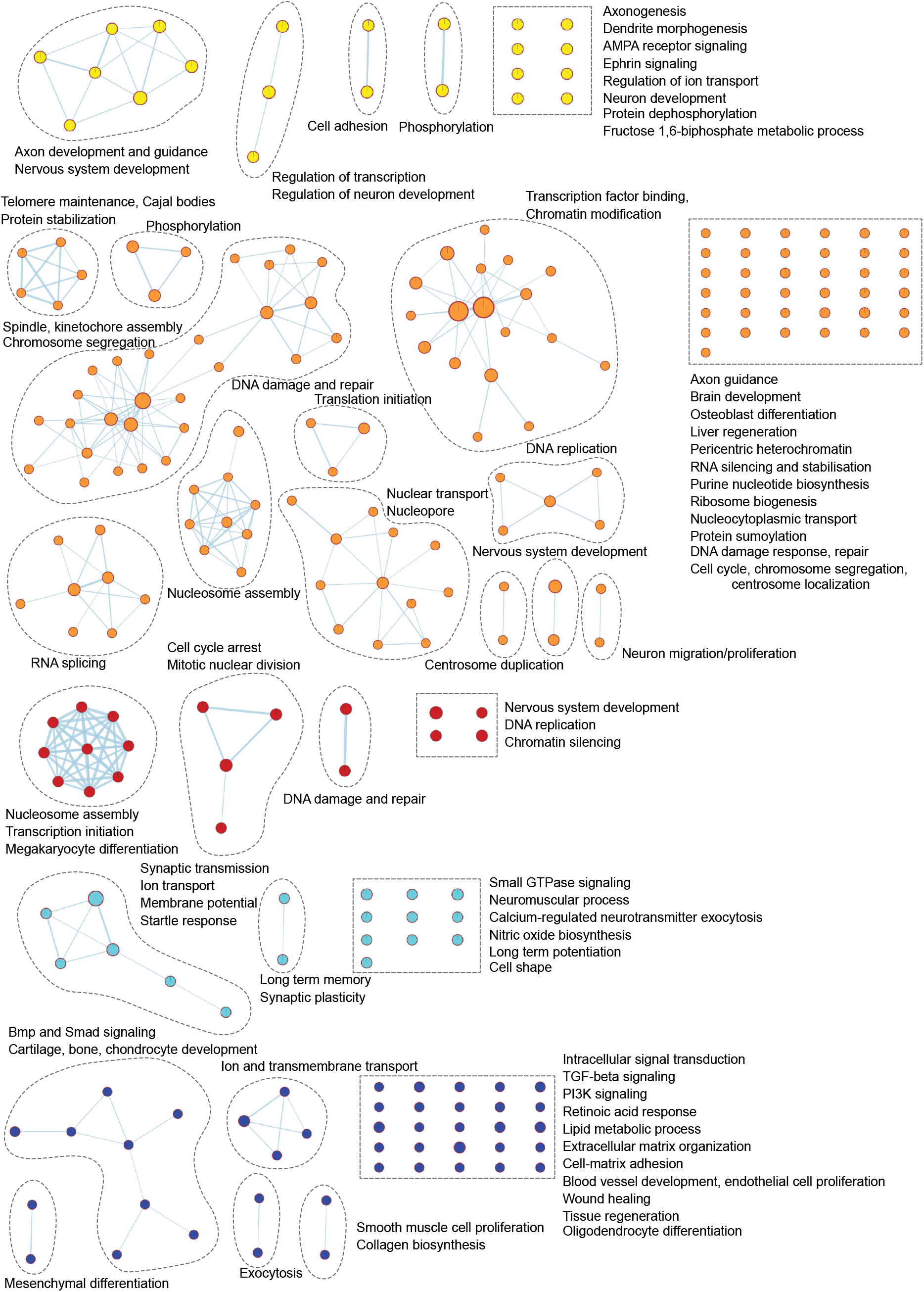
Enrichment of specific biological processes during CGNP development per gene cluster. Gene ontological analysis shows enriched biological processes per gene cluster. Each node represents a biological process. Related biological processes are grouped and labeled by biological theme (curved dashed lines). Individual biological processes are assembled in rectangular boxes (dashed lines). Biological processes connected by edges have genes in common. Enriched biological processes were determined with the Database of Annotation, Visualization and Integrated Discovery (DAVID), v.6.8 (Benjamini-corrected q = 0.1, p =0.01) and visualized with the Enrichment Map app in Cytoscape. Yellow nodes: E15.5 - E17.5 cluster; orange nodes: E15.5 - P7 clusters; red nodes: P0 - P7 cluster; light blue nodes: P14 - P30 cluster; dark blue nodes: P14 - P30 cluster.

**Figure S2 related to Figure 3.**
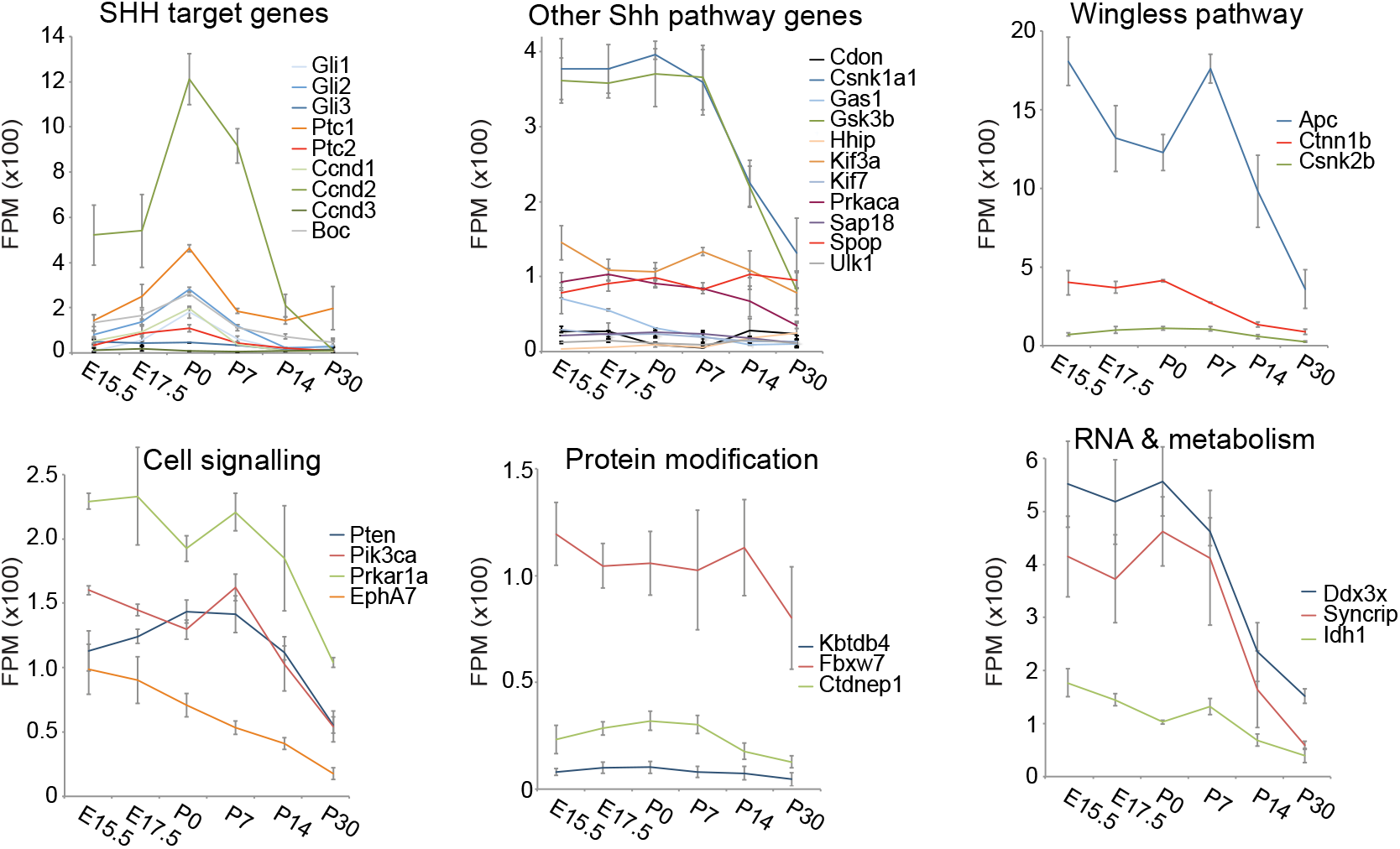
Gene expression and proliferation in developing CGNPs. Gene expression profiles of CGNP genes commonly mutated in medulloblastoma extracted from the RNA-seq data set. Curves represent the average expression level from three biological replicates, error bars indicate standard deviation (FPM=fragments per million).

**Figure S3 related to Figure 6.**
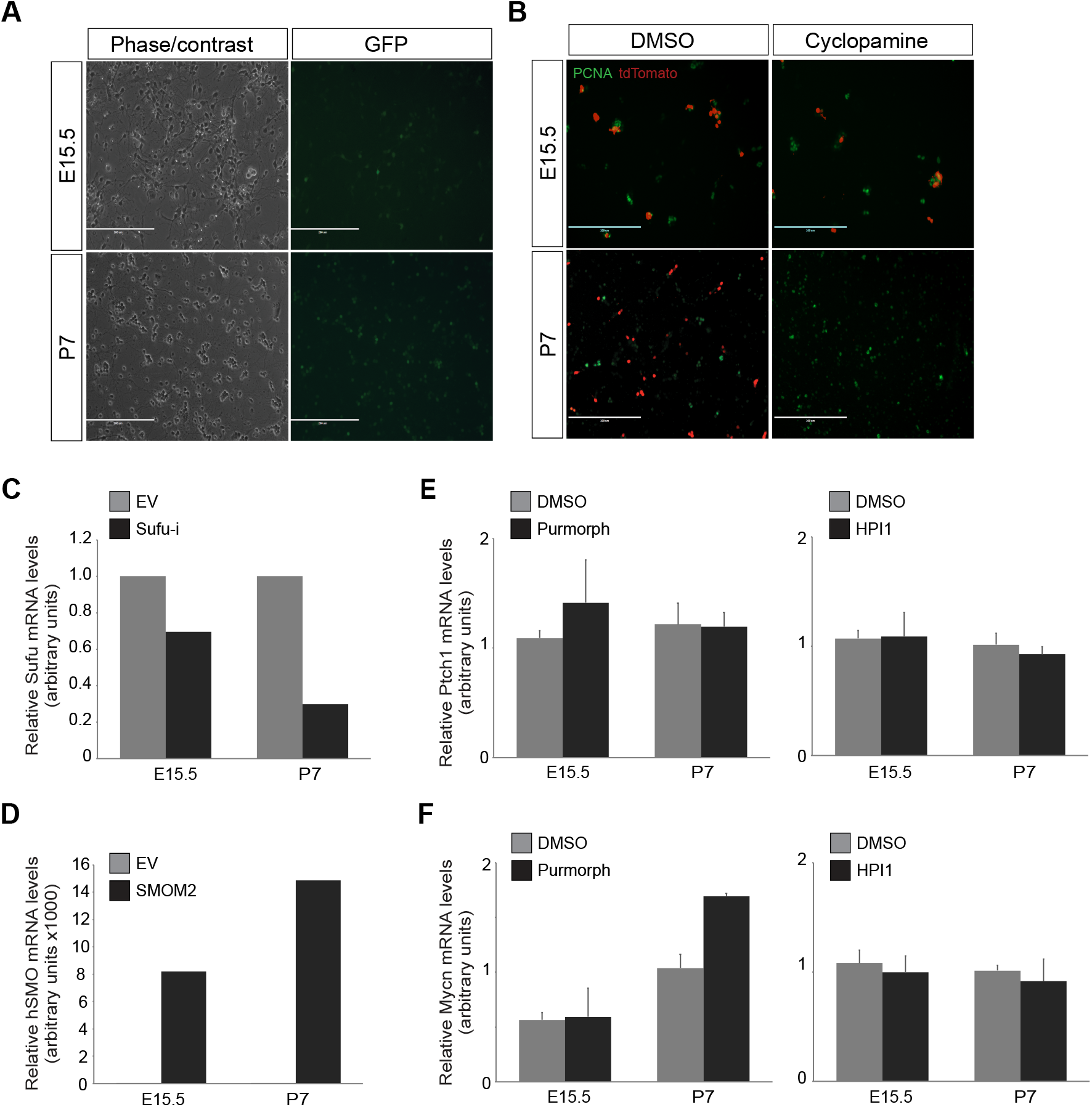
Lentiviral transduction, proliferation and gene expression in primary CGNP cultures. (A) GFP fluorescence (right panels) showing representative lentiviral transduction efficiency in E15.5 and P7 CGNPs. Scale bars indicate size. (B) Representative images of E15.5 and P7 primary CGNP cultures immunolabeled for PCNA and tdTomato after treatment with Cyclopamine or DMSO (mock). (C) qRT-PCR showing relative Sufu mRNA expression levels compared to Gapdh in Sufu shRNA versus empty vector control (EV) transduced E15.5 and P7 CGNPs. (D) qRT-PCR showing relative hSMO mRNA expression levels compared to Gapdh in SmoM2 versus empty vector control (EV) transduced E15.5 and P7 CGNPs. (E-F) qRT-PCR showing relative Ptch (e) or Mycn (f) mRNA expression levels compared to Gapdh in Purmorphamine (left panels) or HPI1 treated (right panels) E15.5 and P7 CGNPs.

**Table S1, related to Figure 1**.

See Excel file.

**Table S2, related to Figure 1**.

See Excel file.

**Table S3, related to Figure 2**.

See Excel file.

**Table S4, related to Figure 3**.

See Excel file.

**Table S5, related to Figure 3**.

See Excel file.

**Table S6.**
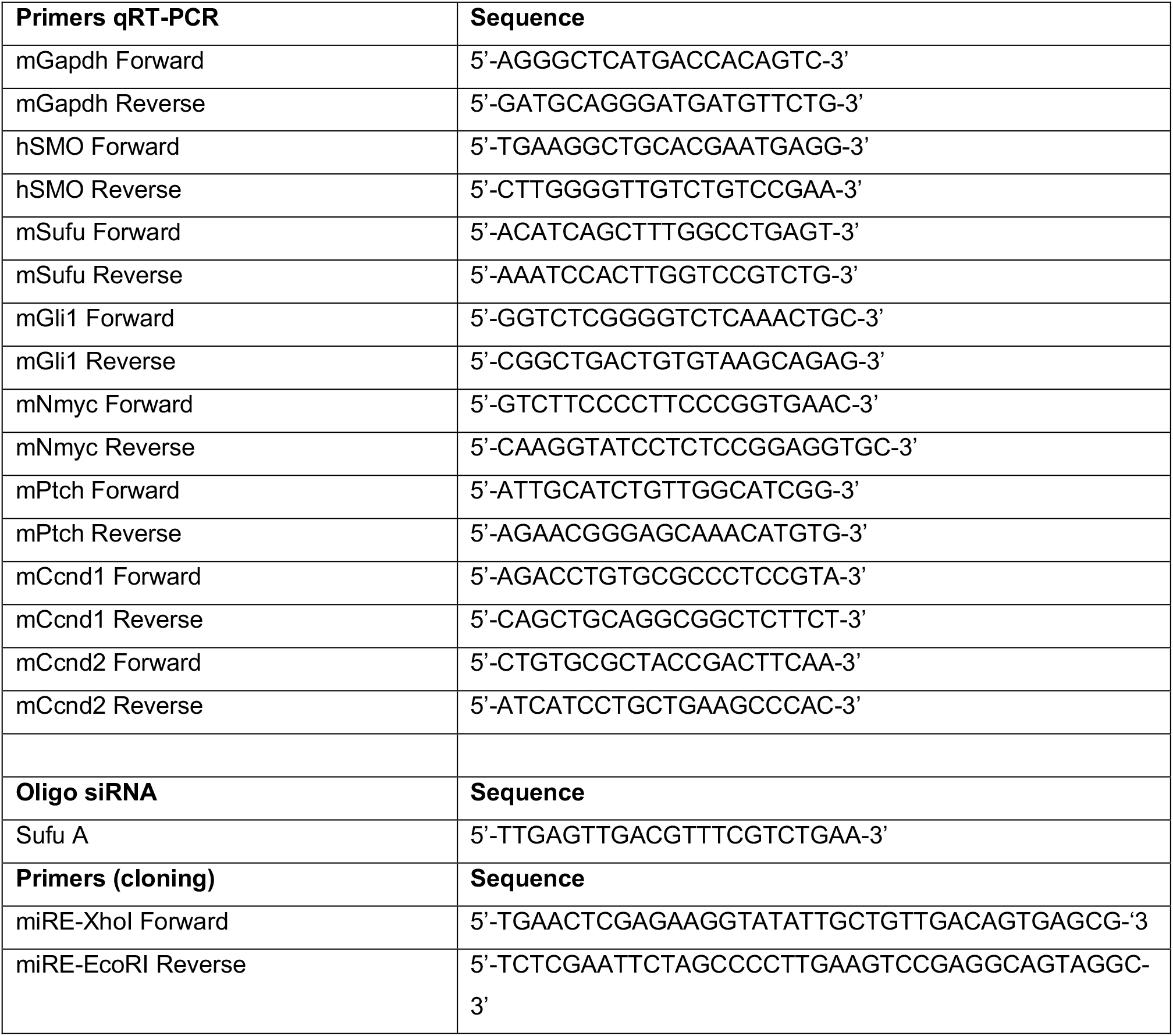
related to Materials and methods.

